# Measuring three-dimensional dynamics of platelet activation using 3-D quantitative phase imaging

**DOI:** 10.1101/827436

**Authors:** SangYun Lee, Seongsoo Jang, YongKeun Park

## Abstract

Platelets, or thrombocytes, are anucleated tiny blood cells with an indispensable contribution to the hemostatic properties of whole blood, detecting injured sites at the surface of blood vessels and forming blood clots. Here, we quantitatively and non-invasively investigated the morphological and biochemical alterations of individual platelets during activation in the absence of exogenous agents by employing 3-D quantitative phase imaging (QPI). By reconstructing 3-D refractive index (RI) tomograms of individual platelets, we investigated alterations in platelet activation before and after the administration of various platelet agonists. Our results showed that while the integrity of collagen-stimulated platelets was preserved despite the existence of a few degranulated platelets with developed pseudopods, platelets stimulated by thrombin or thrombin receptor-activating peptide (TRAP) exhibited significantly lower cellular concentration and dry mass than did resting platelets. Our work provides a means to systematically investigate drug-respondents of individual platelets in a label-free and quantitative manner, and open a new avenue to the study of the activation of platelets.

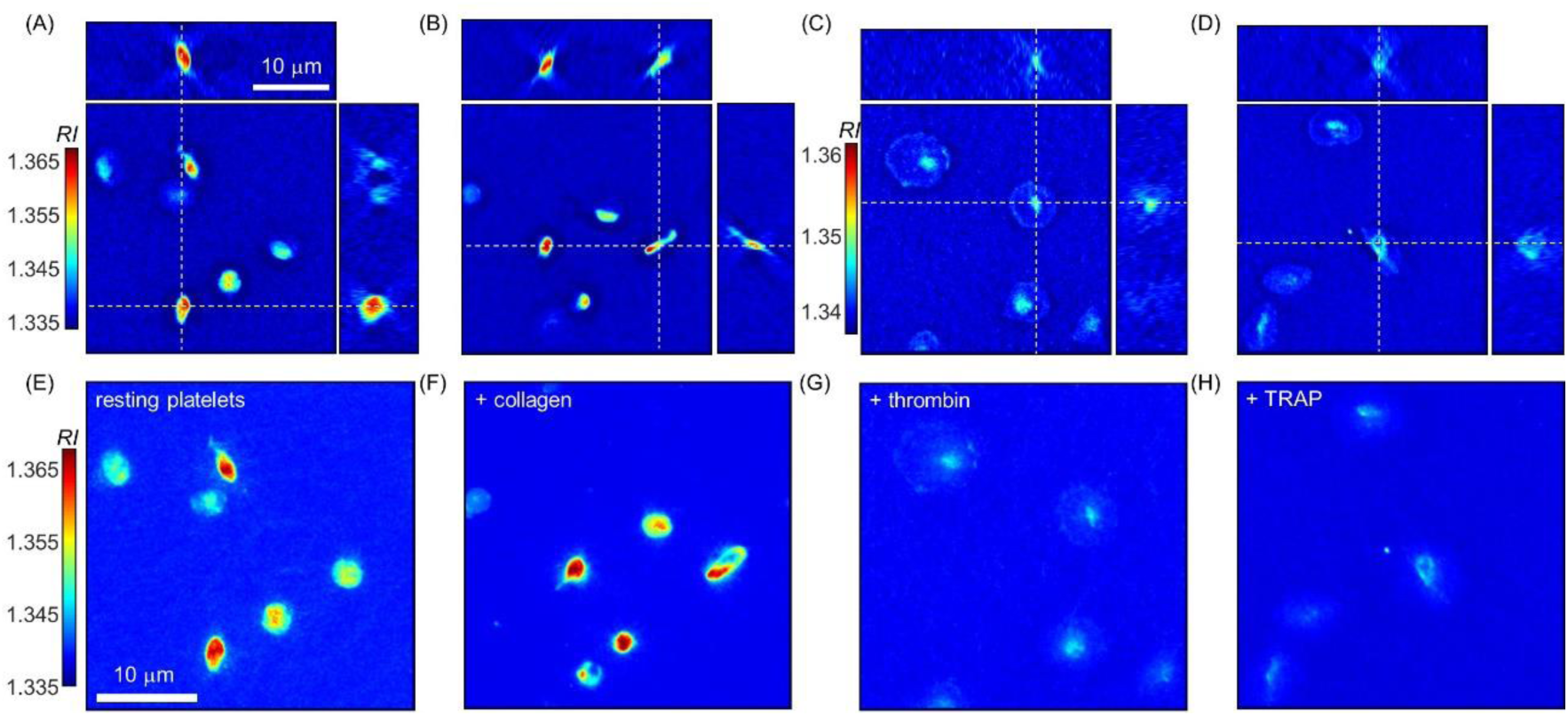

## Introduction

Platelets play an indispensable role in the hemostatic properties of whole blood by detecting injured sites at the surface of blood vessels and forming blood clots^1,2^. More than 10 million people suffer or die annually due to hemostatic failures linked to platelet dysfunctions. One of the most important clinical issues is to determine the appropriate dosage of antiplatelet drugs for treating patients found between hemorrhage and embolism^3-5^. Although blood tests screening for platelet aggregation is performed, the types and dosage of antiplatelet drugs are not determined based on the aggregation test, because controversies regarding their actual efficacy and patient-to-patient variations remain^6-9^ and existing aggregation test mainly address plate aggregation. In extension to these concerns, many researchers have put their efforts into finding clinical clues not only relating to platelet aggregation but also the activation state of individual platelets. However, the particularly small size and atypical shapes of individual platelets have hindered such attempts focused on using biochemical and morphological features of individual platelets for assessing platelet functionality.

The most direct way to access the hemostatic abilities of platelets is to image individual platelets. Various imaging techniques, from electron^10-15^ and scanning probe microscopy^16-19^ to phase-contrast^19,20^ and differential interference contrast microscopy^21,22^ to fluorescence confocal microscopy,^22-24^ have been employed. There is a considerable missing link between the imaging capabilities of electron microscopes that are capable of imaging sub-micron intraplatelet structures but only on dead cells and that of conventional optical microscopes that are capable of imaging live cells but lack of high-resolution quantitative imaging. Atomic force microscopy or scanning ion-conductance microscopy lies in between, but always requires a height reference to precisely reconstruct the topography of live cells, without addressing tomographic internal information^25,26^. Accordingly, only activated platelets stably attached to the substrate can be evaluated. Meanwhile, most platelet analyzers utilized in clinical practice mainly deal with the macroscopic properties of blood plasma and thus cannot fully address the underlying microscopic intraplatelet phenomena.

Here, we present a method to assess the three-dimensional (3-D) morphology and dynamics of individual platelets. We employed quantitative phase imaging (QPI)^27^ and performed measurements of the morphological and biochemical alterations of resting platelets undergoing activation in a label-free, intrinsically quantitative, and real-time manner. We investigated 2 major issues associated with platelet activation: (i) statistical analyses on morphological and biochemical characteristics of resting and agonist-induced activated platelets, and (ii) continuous monitoring of the activation process of individual resting platelets. To selectively activate the respective route of the platelet activation process, either collagen, the more potent thrombin, or thrombin receptor activating peptide (TRAP) were chosen as platelet agonists.

## Results

### 3-D quantitative phase imaging of individual platelets

To measure the 3-D images of individual platelets, we exploited optical diffraction tomography (ODT) (Fig. 1). Due to its high-resolution, label-free, and quantitative imaging capability, ODT has been recently utilized in various studies, including research in the fields of haematology^28,29^, infectious diseases^30,31^, drug discovery^32^, and cell biology^33^. Analog to X-ray computed tomography, ODT reconstructs the 3-D RI of a microscopic object using a laser beam^34-37^. Multiple 2-D hologram images of a sample are measured with various illumination angles, from which the 3-D RI tomogram is reconstructed [Fig. 1(A)]. Here, we utilized off-axis Mach-Zehnder interferometry equipped with a digital micromirror device (DMD)^38,39^ [Fig. 1(B)].

**Figure 1.**
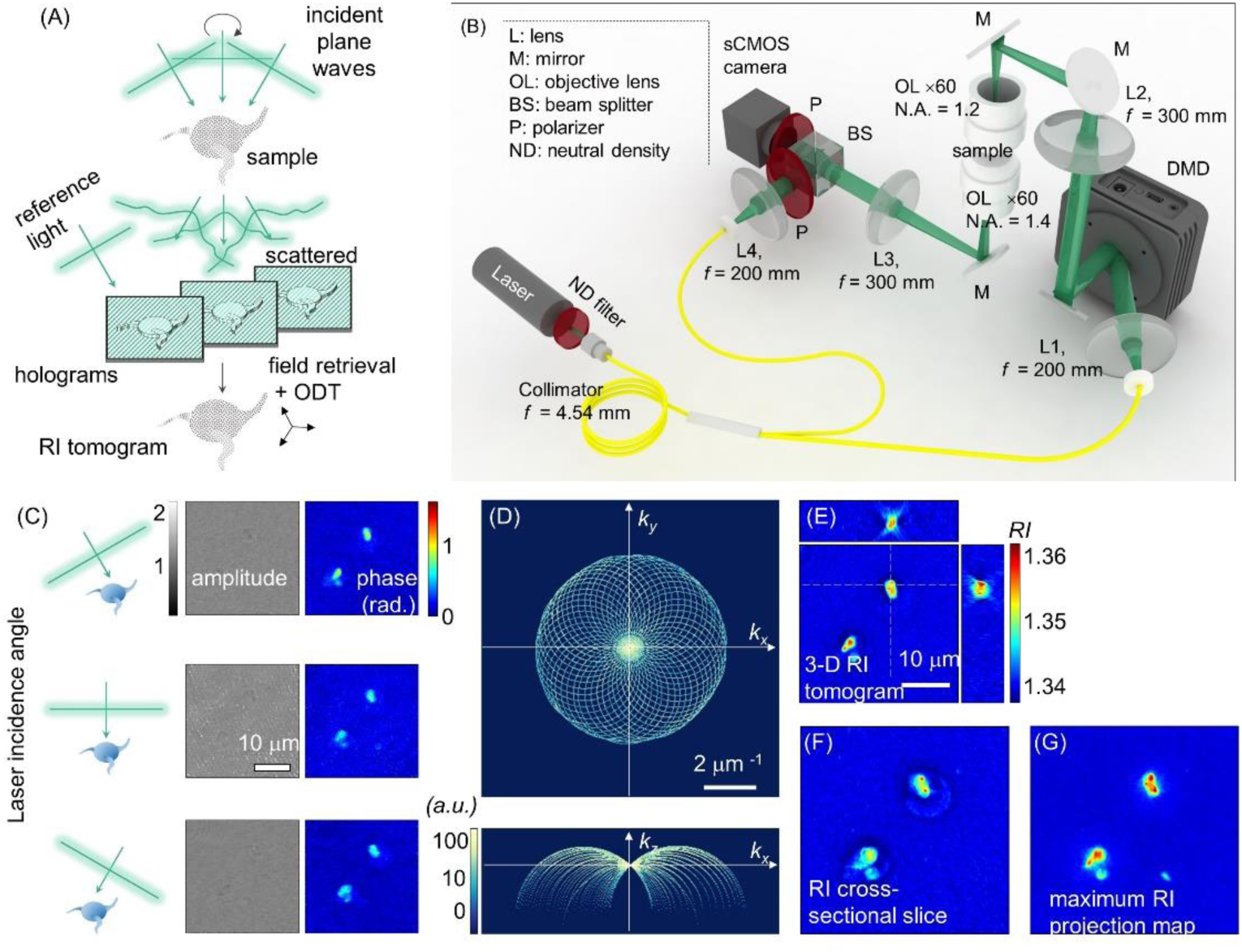
Schematic representation and demonstration of the method. (A) The principle of ODT. (B) The imaging system used. (C) Retrieved 2-D optical fields obtained with various illumination angles. (D) 3-D Fourier spectrum of the synthesised optical scattering potential. (E) Three cross-sectional slices of the reconstructed RI tomograms of platelets. (F) An *x*-*y* cross-sectional slice of the platelet tomogram at *z* = −1.2 μm. (G) 2-D maximum RI projection map of the platelets along the axial direction.

The angle of the laser beam impinging onto the sample was controlled by projecting hologram patterns on the DMD, and the field information of the beam diffracted from the sample was holographically recorded [Fig. 1(C)] (See Supplementary Information). The measured multiple 2D optical field images were mapped in the 3-D Fourier space to synthesize the optical scattering potential [Fig. 1(D)]. Then, the 3-D RI tomogram of the platelet was obtained by taking the inverse Fourier transform on the synthesized scattering potential [Fig. 1(E)]. The cross-sectional image along the *x-y* plane showed a clear spreading of the activated platelets [Fig. 1(F)]. The maximum RI projection (MRP) image of the platelet tomogram, calculated by projecting the maximum RI value of 3-D tomogram along the axial direction, showed the overall shapes of thin cells located at different axial planes at once [Fig. 1(G)].

### Activation of resting platelets by the administration of platelet agonists

To investigate the biochemical and morphological changes of resting platelets during their activation process, we used 3 representative platelet agonists: collagen, thrombin, and thrombin receptor-activating peptide (TRAP) (see *Methods*). First, a collagen-coated coverslip was used to experimentally simulate hemostasis in vivo, in which resting platelets are activated in response to collagen exposed in the wounded blood vessel walls. Then, thrombin (1 U/mL), known as one of the more potent and standard platelet agonists, was used to activate resting platelets more reliably. TRAP is a peptide that interacts with a platelet thrombin receptor activating it in a similar to thrombin manner. Since TRAP does not interact with other plasma proteins, such as fibrinogen, it was used to investigate common morphological and biophysical alterations of resting platelets in an activation process co-stimulated by thrombin. In our experiments, a TRAP solution of 33 μM was used to fully activate resting platelets. Representative 3D RI tomograms of resting platelets and activated platelets stimulated by either collagen, thrombin, or TRAP are presented in Fig. 2. Resting platelets have an elliptical shape with various aspect ratios and mean intraplatelet RI values [Fig. 2(A and E)]. On the other hand, collagen-activated platelets [Fig. 2(B and F)] exhibit more diverse shapes, having pseudopods or intraplatelet regions with inhomogeneous RIs that might be a consequence of an induced degranulation process. Meanwhile, in the case of thrombin- or TRAP-activated platelets, noticeable spreading was observed [Fig. 2(C and D)], thus exhibiting lower RIs than the resting and collagen-stimulated platelets. 20 min after loading of the plasma on the collagen-coated coverslip, most of the platelets were still moving in the plasma and were not stably attached on the surface of the coverslip, as in the case of resting platelets (data not shown). However, thrombin- or TRAP-stimulated platelets were observed to be immobile and were generally spread on the coverslip surface.

**Figure 2.**
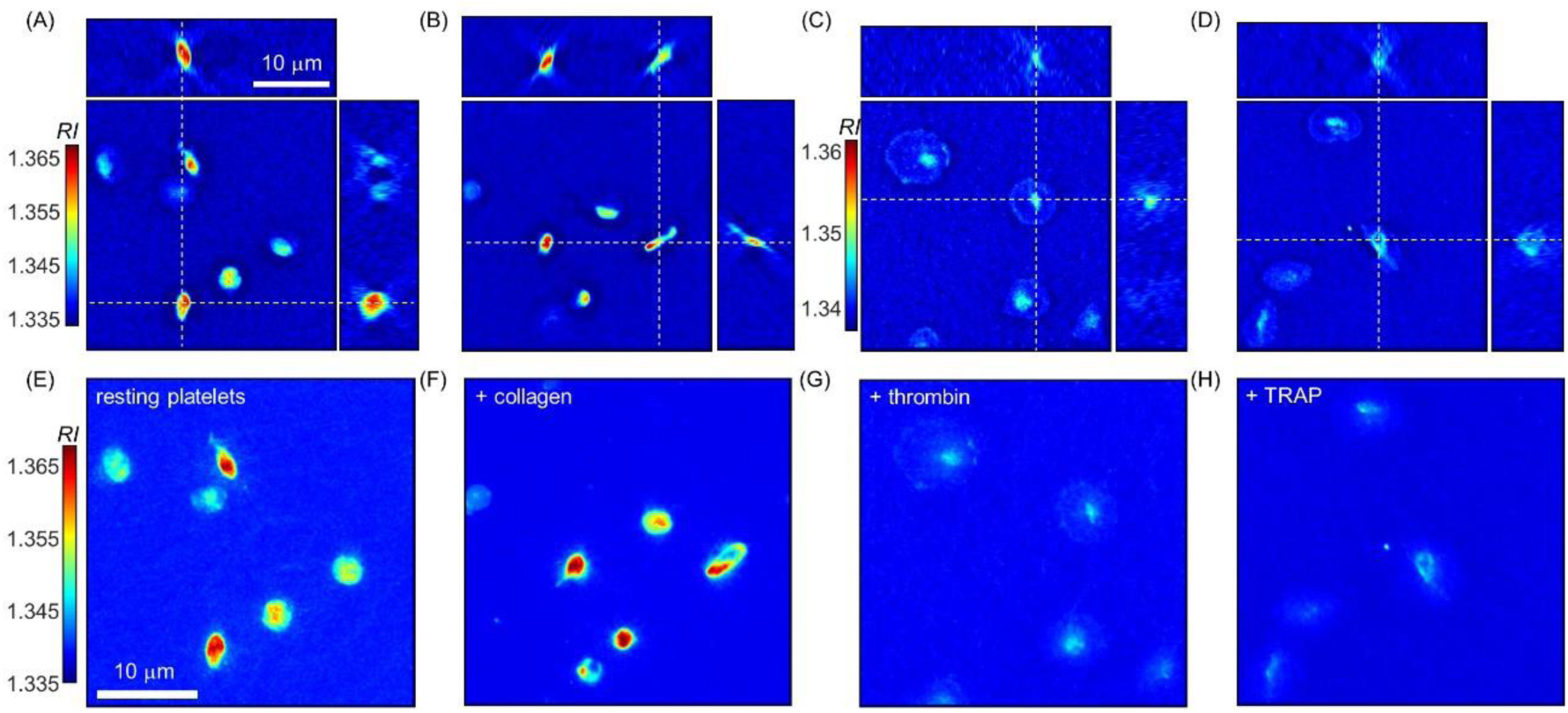
(A–D) Three *x* - *y, y* - *z*, and *z* - *x* cross-sectional slices and (E–H) corresponding MRP images of 3-D RI tomograms of representative resting platelets (A and E) and activated platelets stimulated by either collagen (B and F), thrombin (C and G), or TRAP (D and H).

### Alterations in the biochemical properties of resting platelets during the induced activation process

The biochemical parameters, including mean intraplatelet RI and dry mass of platelets, were retrieved from the measured 3-D RI tomograms (Fig. 3). While there was no statistical difference in the mean intraplatelet RI between resting and collagen-treated platelet groups, the mean RI of the platelet group stimulated by either thrombin or TRAP was significantly lower compared to that of the corresponding control group [Fig. 3(A)-(C)]. Resting and collagen-treated platelet groups had a mean RI of 1.3472 ± 0.0038 and 1.3469 ± 0.0045, respectively. Resting platelets and thrombin-activated platelets exhibited a mean RI of 1.3463 ± 0.0031 and 1.3447 ± 0.0037, respectively [*p*-value < 0.001]. The mean RI of the TRAP-activated platelet group (1.3449 ± 0.0034) was also far lower than that of the control (1.3463 ± 0.0031) [*p* < 0.001]. The presence of some platelets with mean RIs greater than 1.359 in both the collagen- and thrombin-treated platelet groups [red dashed circles, Fig. 3 (A and B)], despite the fact that the mean RI of these platelet groups was equal to or less than that of the respective resting platelet group and there was no resting platelet with mean RI larger than 1.359. However, no such platelets were identified in the TRAP-treated platelet group.

**Figure 3.**
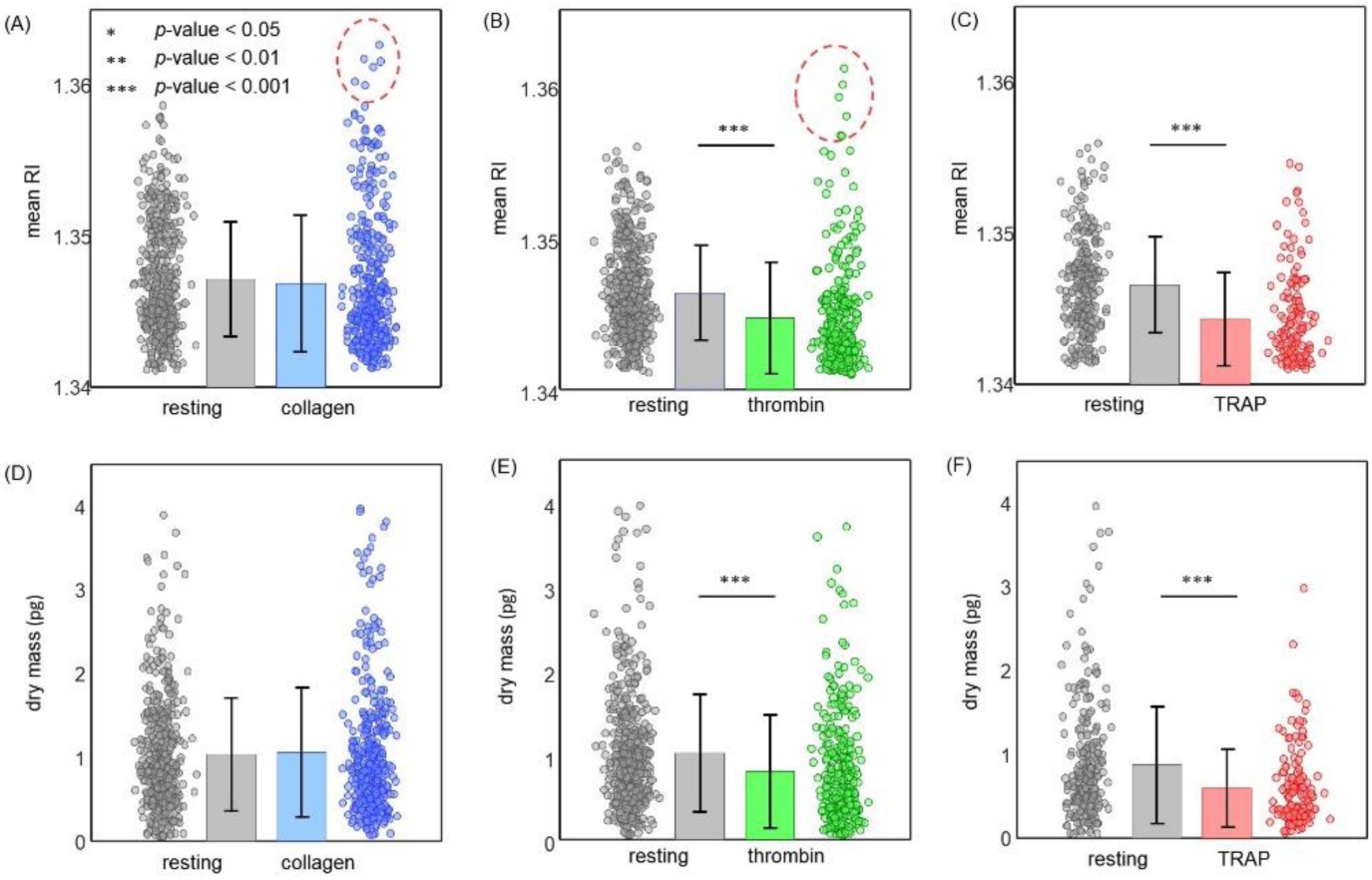
Biochemical properties including mean RI (A–C) and dry mass (D–F) of resting platelets and activated platelets induced by either collagen (A and D), thrombin (B and E), or TRAP (C and F). Each plotted coloured circle denotes a single measurement. Bar height and error bar length indicate sample mean value and standard deviation, respectively. A total of 5, 6, and 3 subjects participated in the experiments using either collagen, thrombin, or TRAP as platelet agonists, respectively. For each participant, more than 50 individual platelets were measured per respective control and agonist-treated platelet groups. The total number of measured resting and collagen-treated platelets were 430 and 356, those of resting and thrombin-treated platelets were 435 and 286, and those of resting and TRAP-treated platelets were 254 and 140, respectively.

Concurrently, the trend observed in the dry mass of agonists-exposed platelets was similar to that of mean RI, implying that the reduced mean RIs of the thrombin- and TRAP-stimulated platelets were mainly caused by a substantial loss of platelet cellular materials, i.e. degranulation, in conjunction with platelet activation. Moreover, this degranulation process during platelet activation might be related to the previously observed suppression of lateral movements observed in thrombin- and TRAP-activated platelets. Specifically, the mean dry mass of resting and collagen-treated platelets was 1.04 ± 0.68 and 1.06 ± 0.77 pg, respectively [Fig. 3(D)]. As expected, the observed difference was not statistically significant. Meanwhile, the mean dry mass of the thrombin-treated platelet group (0.81 ± 0.67 pg) was significantly lower than that of the resting platelet group (1.03 ± 0.70 pg) [Fig. 3(E)], and the mean dry mass of the TRAP-treated platelet group (0.60 ± 0.46 pg) was also significantly lower than that of the resting platelet group (0.87 ± 0.69 pg) [Fig. 3 F)] (both *p*-values < 0.001).

### Alterations in the morphologies of resting platelets during activation process induced by either collagen, thrombin, or TRAP platelet agonists

Figure 4 shows the recorded morphological parameters, including platelet volume, surface area, and sphericity of resting platelets and activated platelets stimulated by either collagen [Figs. 4(A, D, and G)], thrombin [Figs. 4 (B, E, and H)], or TRAP [Figs. 4(C, F, and I)].

**Figure 4.**
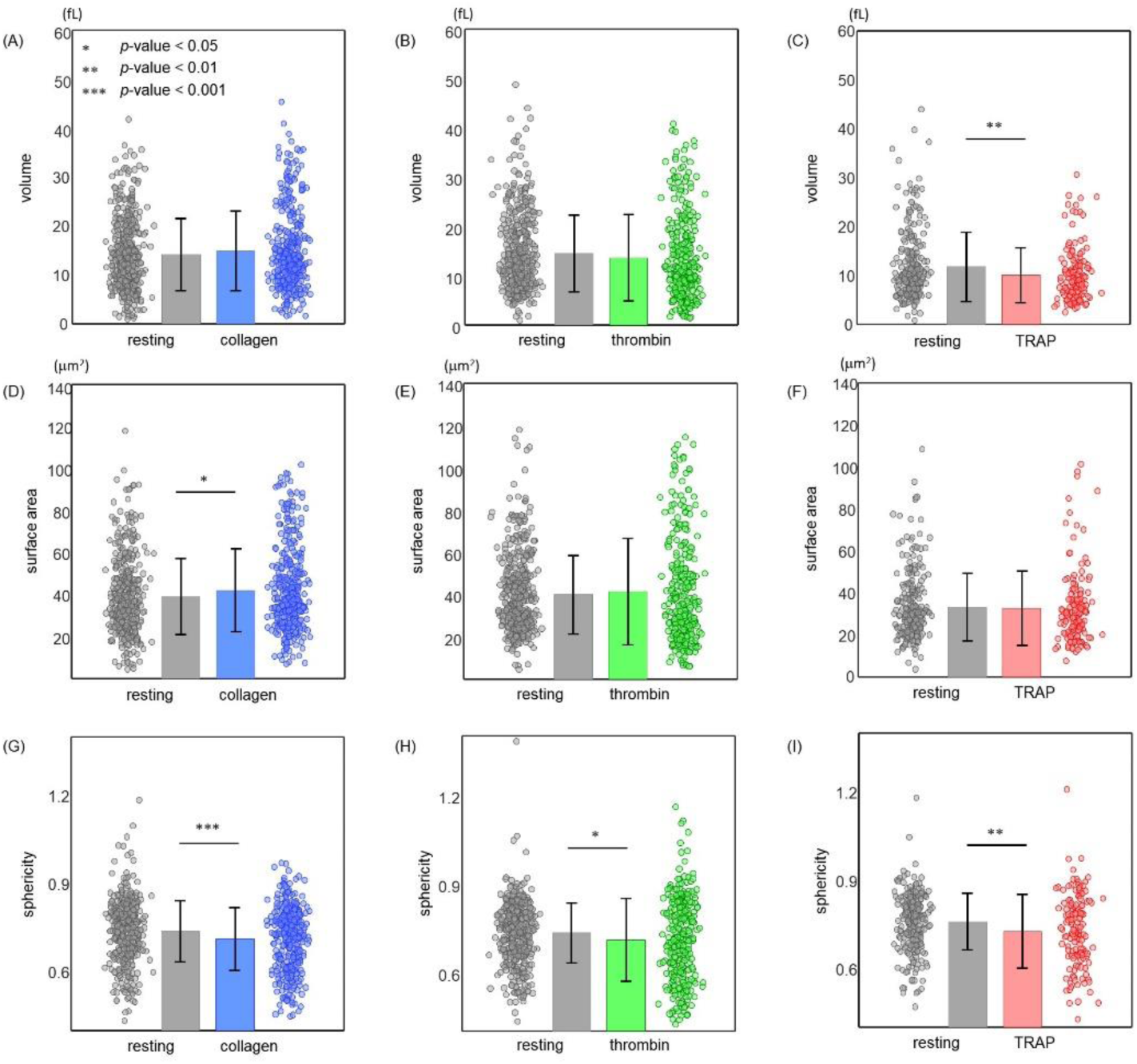
Recorded morphological parameters including platelet volume (A–C), surface area (D–F), and sphericity (G–I) of resting and activated platelets induced by either collagen (A, D, and G), thrombin (B and E, and H), or TRAP (C, F, and I). Each plotted coloured circle denotes a single measurement. Bar height and error bar length indicate sample mean value and standard deviation, respectively.

The resting platelet group displayed a mean volume of 14.2 ± 7.4 fL, slightly below the mean volume of 14.9 ± 8.1 fL shown by the collagen-treated platelet group, and thus not statistically significant [Fig. 4(A)]. Contrary, in the case of thrombin-treated platelets, the mean volume of resting platelets (14.5 ± 7.8 fL) was slightly but not significantly higher than that of the thrombin-activated platelets (13.6 ± 8.8 fL) [Fig. 4(B)]. However, TRAP-activated platelets exhibited a significantly lower mean cellular volume (10.1 ± 5.6 fL) compared to that of the resting platelet group (11.8 ± 7.0 fL) [*p* < 0.01; Fig. 4(C)].

While the collagen-treated platelet group showed a higher mean surface area compared to the resting platelet group, there was no statistical difference in the mean surface area between the thrombin- or TRAP-treated platelet group and the corresponding control group. The mean surface area of collagen-treated platelets (42.1 ± 19.9 μm^2^) was statistically higher than that of the resting platelets (39.1 ± 18.0 μm^2^) [*p* = 0.03; Fig. 4(D)]. However, resting and thrombin-activated platelets had a mean surface area of 40.0 ± 18.6 and 41.3 ± 25.1 μm^2^, respectively (*p*-value = 0.43; Fig. 4(E)). Similarly, the mean surface area of the resting and TRAP-treated platelet groups was 33.2 ± 16.0 and 32.6 ± 17.8 μm^2^, respectively [*p*-value ∼ 0.73; Fig. 4(F)].

The most prominent alteration in the morphology of resting platelets following administration of platelet agonists was observed in platelet sphericity, a parameter measuring how closely a cell resembles a sphere [Fig. 4(G–I)] The mean sphericity of the agonist-activated platelet groups was significantly lower than that of the respective control group. Resting and collagen-activated platelets had a mean sphericity of 0.74 ± 0.10 and 0.71 ± 0.11, respectively (*p* < 0.001). In the case of thrombin treatment, both the slightly reduced volume and slightly increased surface area of the thrombin-activated platelet group, although not statistically significant, resulted in a statistically significant (*p*-value ∼ 0.01) lower mean sphericity (0.71 ± 0.14) compared to that of the resting platelet group (0.73 ± 0.10). Again, the mean sphericity of the TRAP-treated platelet group (0.73 ± 0.13) was significantly lower than that of the control group (0.76 ± 0.10), mainly resulting from the reduced mean volume of the TRAP-activated platelets (*p*-value ∼ 0.006). Detailed information on the individual participants of these experiments are presented in Fig. 5.

**Figure 5.**
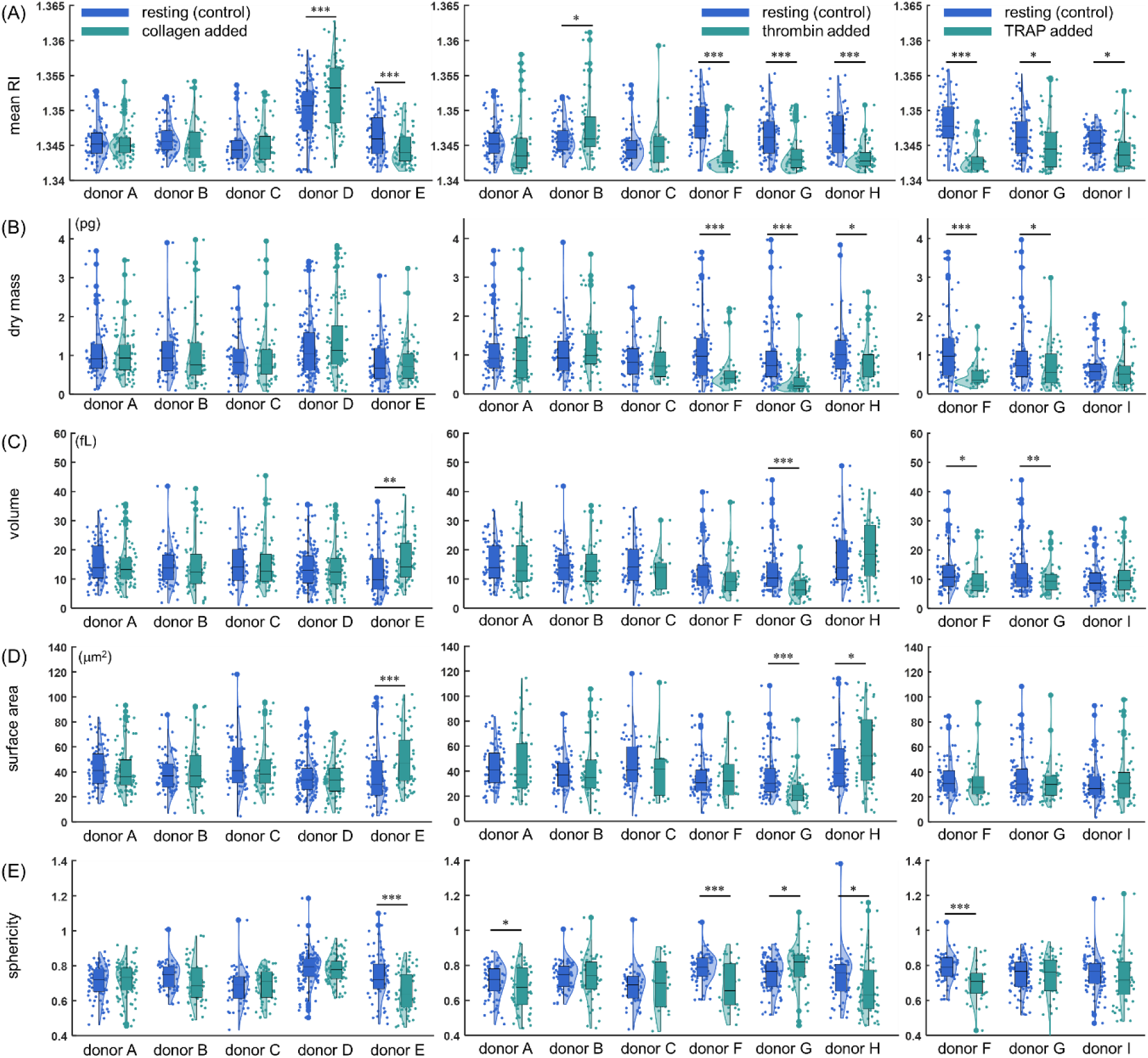
The biochemical and morphological platelet parameters of individual subjects including (A) mean RI, (B) dry mass, (C) volume, (D) surface area, and (E) sphericity. Each colored circle in the plots denotes single measurement. The box plots present mean values with upper and lower quartiles. The vertical line of box plots indicates a parameter range of the population. The normalized probability density function on the measured platelet parameters of individual donor is also specified.

### Continuous monitoring of individual platelets under activation process

We performed the continuous optical measurements on individual platelets during thrombin-induced activation. After administration of 1 U/mL thrombin, the entire activation process of each platelet was monitored by obtaining one 3-D RI tomogram every 6 s for a total of 10 min. From the time platelets entered the field-of-view of the microscope and began to attach to the coverslip, various cellular parameters were measured and analyzed [Fig. 6]. Figure 6(A) presents the timely evolution of the mean RI of 10 measured platelets along the whole activation process. The mean RI of all measured platelets was significantly reduced during the activation process induced by thrombin. The mean RI of measured platelets at the beginning and end of the monitored activation process were 1.3446 ± 0.0018 and 1.3428 + 0.0012, respectively (*p* < 0.001). The timely progression of the sphericity of 11 platelets is shown in Fig. 6(B). A paired *t*-test indicated that the mean sphericity of the monitored platelets at the end of the activation process (0.87 ± 0.19) was significantly lower than that of corresponding platelets at the moment of thrombin administration (0.96 ± 0.11) (*p*-value ∼ 0.05). Figure 6(C and D) represents the timely evolution of the volume of a total of 10 platelets. Mean volumes of the monitored platelets during the first 4 and last 4 time-points were 19.7 ± 13.3 and 19.7 ± 17.7 fL, respectively.

**Figure 6.**
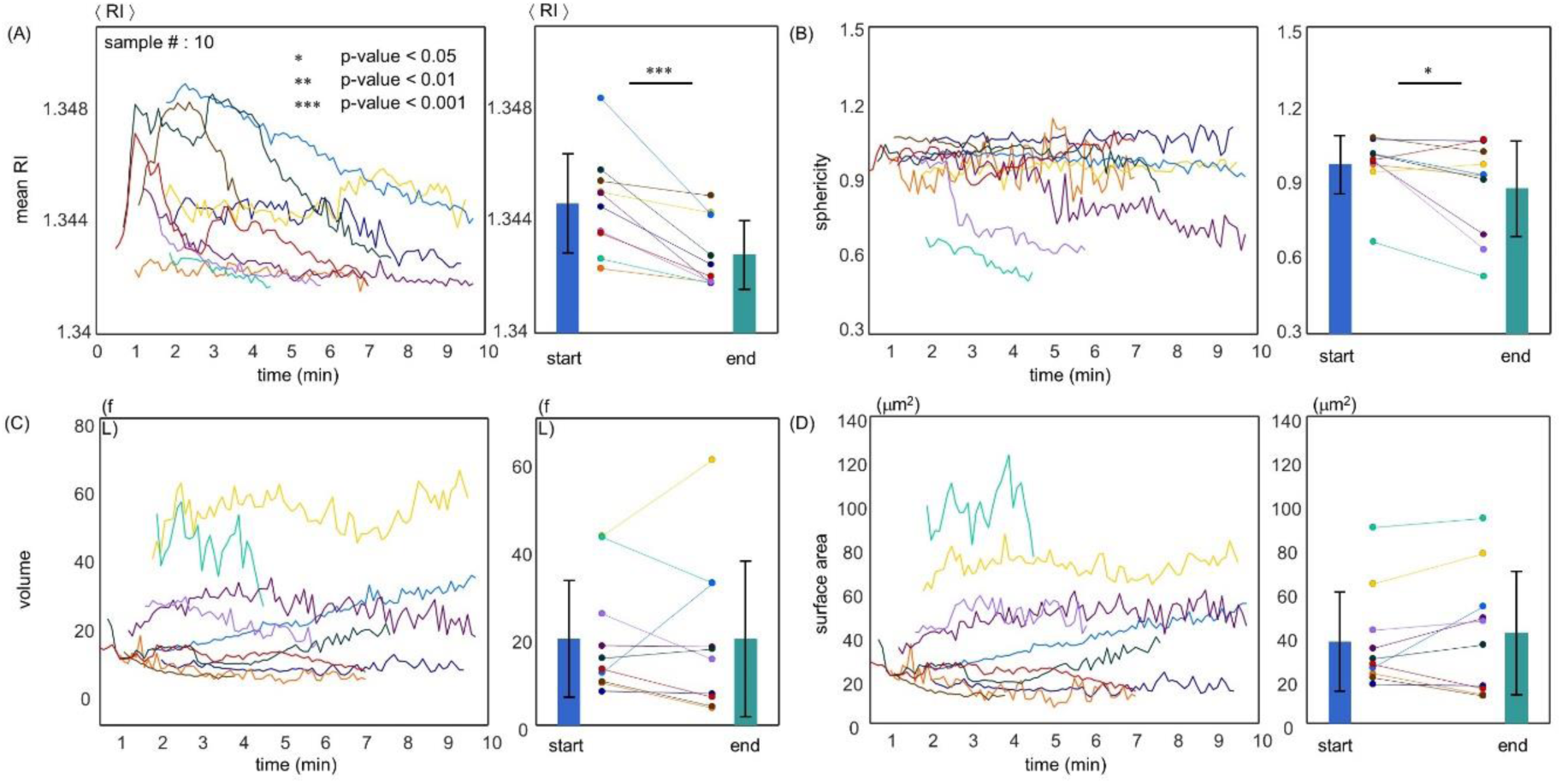
Continuous alterations in mean RI (A), sphericity (B), volume (C), and surface area (D) of individual platelets over both long (10 min, one tomogram per 6 s) time-scale monitoring after administration of thrombin (1 U/mL). Each coloured line represents the time trajectory of the recorded parameter of an individual platelet. The graph on the right of each panel presents the mean first 4 and last 4 recorded time-points for the respective parameter of each individual platelet. Bar height and error bar length indicate sample mean value and standard deviation, respectively.

To further investigate the dynamics of platelets when cell spreading occurred, we repeated the same procedure on a total of 11 individual platelets during activation by 1 U/mL thrombin at the temporal rate of one tomogram per second during the first 100 s of the activation process (Fig. 7). After thrombin treatment, we observed that the mean RI of 6 of the 11 platelets was significantly reduced from a certain time-point onwards, and this overall trend was similar to the results obtained from the long-scale optical measurement [10 min; Fig. 7(A)]. Meanwhile, the remaining 5 platelets did not undergo significant changes in RI during the early stages of activation. The mean RI of the monitored 11 platelets during the first 4 time-points (1.3471 ± 0.0030) was significantly higher than that of the corresponding platelets during the last 4 time-points (1.3447 ± 0.0016) (*p* = 0.01).

**Figure 7.**
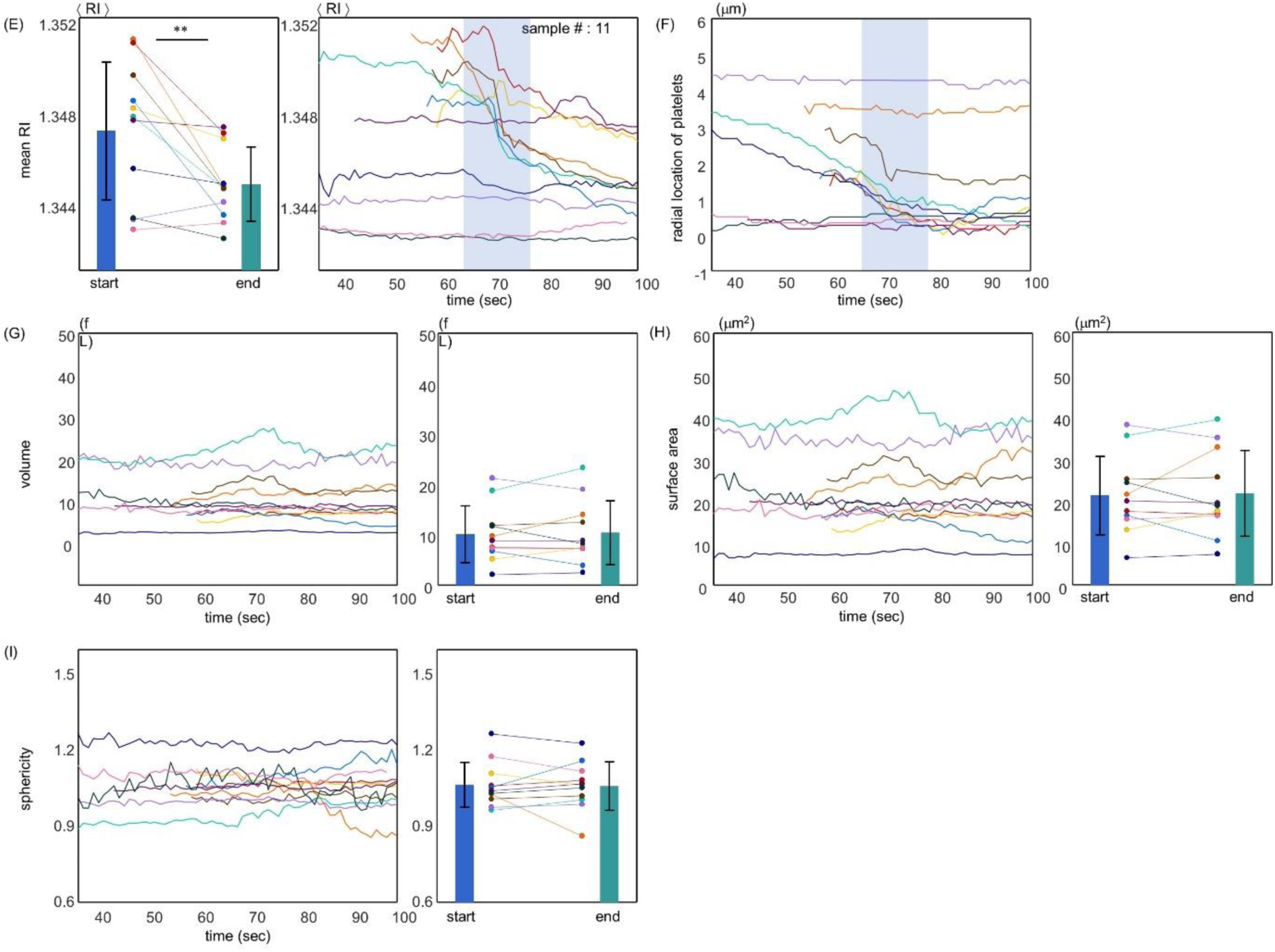
Continuous alterations in mean RI (E), radial coordinates (F), volume (G), surface area (H), and sphericity (I) of individual platelets over short (100 s, one tomogram per s) time-scale monitoring after administration of thrombin (1 U/mL). Each coloured line represents the time trajectory of the recorded parameter of an individual platelet. The graph on the right of each panel presents the mean first 4 and last 4 recorded time-points for the respective parameter of each individual platelet. Bar height and error bar length indicate sample mean value and standard deviation, respectively.

Figure 7(B) represents the recorded radial location of the examined platelets during the activation process, and therefore each slope curve indicates the instant speed of the respective platelet. The timely progression of volume and surface area of all 11 platelets are shown in Fig. 7(C)-(E). The mean volume and surface area of platelets at the beginning of the activation process were 10.1 ± 5.6 fL and 21.1 ± 9.3 μm^2^, whereas those at the end of the optical measurements were 10.4 ± 6.3 fL and 21.7 ± 10.2 μm^2^, respectively. Consecutively, the mean sphericity of the platelets displayed a value of 1.06 ± 0.09 in the initial and a value of 1.06 ± 0.10 in the last stage of the optical measurements. In this instance, the paired *t*-test did not verify the hypothesis that the volume, surface area, and sphericity of resting platelets were significantly altered during the first 100 s after thrombin administration (1 U/mL).

The experiments based on individual cell tracking using a high-speed QPI system also gave similar results. While the sphericity of individual platelets did not change in a statistically significant manner during the first 100 s after thrombin treatment [Fig. 7(E)], it was significantly reduced in the experiments observing the activation process for 10 min [Fig. 6(B)]. This result implied that resting platelets were slowly and gradually flattened as they spread over a coverslip during the whole 10 min of the activation process induced by thrombin. It can be assumed from Fig. 7(B) that 6 of the 11 platelets were initially moving at a constant speed and then stopped at a specific time-point at which platelets adhered to the coverslip. In this case, we observed that the time-point at which platelets stopped moving qualitatively coincided with the time at which the mean RIs of the platelets were significantly decreased. These regions are shaded in light blue in Fig. 7(A and B) to assist visualization. This observation further supported our hypothesis that the process of intraplatelet degranulation is essential for platelets spreading over a substrate.

## Discussion

We found that collagen-stimulated platelets did not exhibit statistically significant differences in the mean intraplatelet RI and dry mass when compared with resting platelets [Fig. 3(A and D)]. This indicated that, despite the existence of some degranulated platelets with a few pseudopods, the given amount of collagen was not sufficient to thoroughly activate all resting platelets and thereby the integrity of a large portion of platelets was maintained. On top of that, most collagen-treated platelets were mobile and moved inside the plasma similar to resting platelets. However, platelets stimulated by a more potent agonist, such as thrombin or TRAP exhibited a significantly decreased mean RI and dry mass relative to those of resting platelets and no longer moved on the coverslip (Fig. 4). This trend was also supported by the result obtained by continuous monitoring of the activation process of individual platelets by 1 U/mL thrombin administration, in both the slow and fast measurements [Fig. 5].

The observed suppression in lateral movements of platelets exposed to thrombin and TRAP with significantly decreased intraplatelet RIs suggested the possibility that a significant loss of platelet cellular materials via degranulation might have preceded the spreading of platelets over the substrates during the process of platelet activation. From the results obtained by the rapid monitoring of the activation process of individual platelets induced by thrombin (one tomogram per s), we found that the time-point at which the RI of the platelet started to decrease, qualitatively coincided with the very timing at which velocity of the platelet was significantly decreased and platelet adhered to the coverslip [Fig. 7(A and B)]. This observation further strengthened our hypothesis that a substantial amount of granular exocytosis should be a prerequisite for platelet spreading. This idea is also consistent with the results of recent research using fluorescent antibodies^40^.

One of the interesting results was that the mean RI of about 30 % of platelets did not monotonically decrease after exposure to thrombin, but to the contrary, slightly increased in the early stages of activation [Fig. 6(A); see also Supplementary Figure 1]. This observation further explained why some platelets in the collagen- or thrombin-treated platelet group, appeared to have a mean RI of larger than 1.359, although mean RI of these platelet groups was equal to or less than the mean RI of resting platelets, which was never greater than 1.359 [Fig. 3(A and B)]. In addition, this interpretation was also consistent with the main result of a previous study showing that intraplatelet RIs increase for a short period of time immediately after the administration of platelet agonists^41^. Also, statistical analysis for the results in Fig. 5 did not verify the hypothesis that activation by thrombin significantly alters platelet volume. This was mainly because individual differences were far too large, consistent with our previous results using statistical approaches in Fig. 3(B).

In the experiments based on statistical approaches, all platelet groups treated with platelet agonists were observed to have a statistically lower mean sphericity than the corresponding control groups, regardless of the particular type of agonists [Fig. 4(G–I)]. For example, in the case of collagen administration, the observed reduced cell sphericity was mainly due to the increased surface area of the platelets. This observation was consistent with previous research reporting that both the formation of pseudopods and spreading over a substrate were achieved by platelets as the invaginated membranes of the platelet open canalicular system (OCS) started to protrude out of the cell. On the other hand, in the case of TRAP administration, the observed reduced cell sphericity was mainly the result of the decreased platelet volume, and this might have been a direct consequence of the continuous RI diminution of platelets through degranulation during the activation process. The results obtained for the thrombin-treated platelet group were thought to reside halfway between the cases of collagen and TRAP administration.

Neither long or short time-scale experiment could verify the hypothesis that thrombin significantly changes the volume of resting platelets [Figs. 6(C) and (G)], and these observations also agreed well with our statistical approach-based results [Fig. 4(E)]. It was noteworthy that the 2 platelets, with increased volumes during activation [Fig. 6(C)], were the platelets with the greatest mean RIs at the end of the 10 min observation [Fig. 6(A)]. Considering that administration of 30 μM TRAP significantly reduced not only the mean RI but also the volume of resting platelets, and our observation that the above 2 platelets were still in the process of decreasing their intraplatelet RIs, we think that administration of more potent agonists or longer observation would eventually lead to a significant reduction in the volume of these platelets via the continuous degranulation process.

One of the most important clinical issues in platelets is to find the sweet spot between the bleeding tendency and the blood clotting of the patient, and thus continuous attempts to make a breakthrough in this field by focusing on the activation process of individual platelets have been tried.

In this study, we presented QPI as an efficient and effective optical imaging tool for assessing various morphological and biochemical properties of individual platelets with quantitative and live-cell imaging capabilities. We characterized the activation process of individual platelets using both statistical and single-cell based approaches. Our results show that the decreasing internal RIs and slightly elevated sphericity of human platelets were the most representative biophysical markers for distinguishing the activation state of individual platelets. We expect that the present method sheds light on the field of testing platelet functionality by providing information about the activation process of individual platelets based on morphology and intraplatelet protein contents.

## Methods

### Sample preparation

Blood was gently drawn from healthy volunteers who had not been subjected to any anticoagulant treatments for at least 2 weeks by standard venepuncture into sodium citrate vacutainers (2.7 mL, 3.2 % v/v sodium citrate; BD, NJ, USA). The informed consent was provided by each volunteer. Within one hour after collection, blood was centrifuged for 12 min at 200 *g* and at room temperature (24 °C) to obtain platelet-rich plasma (PRP)^42,43^. Platelet-poor plasma (PPP) samples were simultaneously obtained by centrifugation at 800 *g* for 15 min for the purpose of platelet dilution. Obtained PRP was then diluted 3 times with PPP to secure 2 or 3 platelets on average in the FOV of the imaging system. Before loading to the microscope stage, diluted platelet suspensions were sandwiched between 2 coverslips (24 × 50 mm; Matsunami, Japan) and sealed to prevent them from drying out. All the above blood gathering protocols were performed at the Pappalardo Medical Center and were approved by the ethics committee of Korea Advanced Institute of Science and Technology (KAIST; Daejeon, Republic of Korea) (IRB project: #KH2017-04).

## Acknowledgments

This work was supported by KAIST, BK21+ program, Tomocube, and National Research Foundation of Korea (2017M3C1A3013923, 2015R1A3A2066550, 2018K000396).

## Authorship contributions

S.L., S.J., and Y.P. conceived the initial idea. S.L. performed the experiments and analysed the data. All authors wrote and revised the manuscript. Correspondence and requests for materials should be addressed to Y.P.

